# Two incursions, two viruses: Emergence of a second novel Shamonda virus clade in Central Europe, 2026

**DOI:** 10.64898/2026.09.05.749452

**Authors:** Kerstin Wernike, Florian Pfaff, Sophie Zeiske, Dirk Höper, Martin Beer

## Abstract

Following the emergence of Shamonda virus (SHAV) in Europe in 2026, we identified a second, genetically distinct SHAV clade in cattle in Germany. Consistent differences across all three genome segments together with divergent regional distribution patterns indicate independent introductions. We have designated the two clades as SHAV Europe 1 (SHAV-EU1) and SHAV Europe 2 (SHAV-EU2). The unexpected co-circulation of these two clades has important implications for diagnostics, surveillance, host range assessment, risk evaluation, and control measures.

## Main text

### Description of the current event

In summer 2026, Shamonda virus (Family: *Peribunyaviridae*; Species: *Orthobunyavirus schmallenbergense*; SHAV), an orthobunyavirus of the Simbu serogroup, emerged in Europe where it was detected in cattle in Switzerland, southern Germany, and France (1, 2). The infections were associated with clinical disease, including fever, diarrhoea and marked decreases in milk production. During the diagnostics of the newly introduced virus in regions of western Germany, that had not yet been affected, an unexpected RT-qPCR reaction pattern was observed. Metagenomic diagnostics and sequence analysis was performed to identify the underlying cause.

### Diagnostic Workflow and Unexpected RT-qPCR Pattern

Following the initial detection of a European SHAV variant (SHAV-EU1) in southern Germany (1), and in the absence of a specific RT-qPCR at that time, a diagnostic workflow was established to distinguish SHAV-EU1 from Schmallenberg virus (Species: *Orthobunyavirus schmallenbergense*; SBV), the only Simbu serogroup virus circulating in Central Europe at that time. The workflow consisted of a generic RT-qPCR for Simbu serogroup viruses (pan-Simbu) (3) followed by an S-segment based specific RT-qPCR for SBV (SBV-S RT-qPCR) (4). SHAV-EU1 tested positive in the pan-Simbu and negative in the SBV-S RT-qPCR (1). Initially, samples from the SHAV-EU1 affected federal state of Baden-Württemberg as well as from the neighbouring states of Bavaria, Rhineland-Palatinate and Hesse reacted as expected. However, suspect samples sent to the German National Reference Laboratory (NRL) for SBV from the western and northern federal states of Lower Saxony and North Rhine-Westphalia in mid-August unexpectedly tested positive in the SBV-S RT-qPCR. For most samples both RT-qPCRs showed comparable quantification cycle (Cq) values (**Supplemental Table S1**). To corroborate the SBV-related results, L- and M-segment based SBV RT-qPCRs were also performed (5), but tested negative. The same combination of positive pan-Simbu and positive SBV-S RT-qPCR was observed in a subset of samples from Hesse, where previously only SHAV-EU1 had been detected. Therefore, in Hesse we detected a mixture of RT-qPCR reaction patterns. The date of the first sample submission to the NRL for SBV per state and the detected SHAV clade is shown in **Figure 1**.

**Figure 1:**
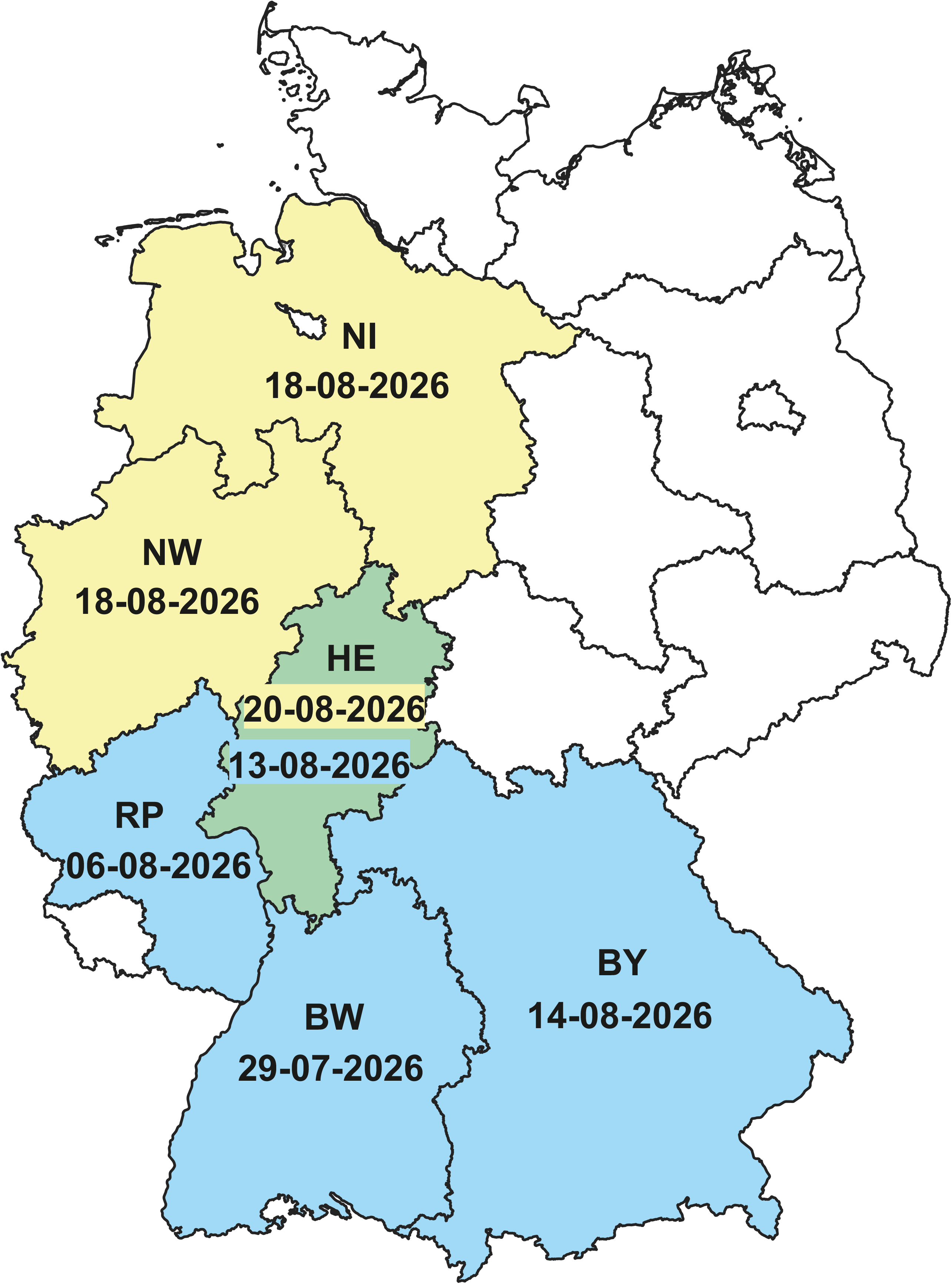
First detections and spatial distribution of the newly introduced Shamonda virus clades in Germany. The dates at which the samples were received at the German National Reference Laboratory for Schmallenberg virus, Friedrich-Loeffler-Institut, are given. Regions where Shamonda virus Europe 1 (SHAV-EU1) has been detected are highlighted in blue, where Shamonda virus Europe 2 (SHAV-EU-2) is present in yellow and the federal state where both clades have been found is shown in green. BW - Baden-Württemberg, BY - Bavaria, HE - Hesse, NI - Lower Saxony, NW - North Rhine-Westphalia, RP - Rhineland-Palatinate.

To determine the underlying cause of the unexpected RT-qPCR pattern, a selection of representative samples was analysed by metagenomic sequencing (Supplemental Material). For all selected samples, a segmented viral genome with similarity to orthobunyaviruses was obtained.

Phylogenetic analyses were performed separately for coding regions of all three genome segments (Supplementary Material).

### A Divergent Shamonda Virus Clade from Northwestern Germany

The virus identified from the samples with the unusual RT-qPCR pattern was classified as SHAV based on phylogenetic analysis of the amino acid sequences of the L protein, the glycoprotein precursor and the nucleocapsid protein (**Supplemental Figures S1-3**). This was also supported by pairwise amino acid identities, as sequences of the European SHAV variant detected in 2026 (SHAV-EU1) and the Nigerian SHAV strain IbAn5550 consistently showed the closest relationship to the newly sequenced virus across all three genome segments (**Supplemental Figure S4**). However, the newly identified SHAV was not identical to the previously reported European SHAV-EU1 clade.

Phylogenetic analyses of the nucleotide sequences of the coding regions of the L protein, the glycoprotein precursor and the nucleocapsid protein were conducted using all available SHAV sequences and appropriate outgroups when possible. Interestingly, the novel European SHAV sequences did not cluster with SHAV-EU1 (**Figure 2A, B, and C**) but instead formed a distinct phylogenetic group across all three genome segments. The most closely related sequences were from SHAV strains previously detected in Japan, with mean pairwise nucleotide identities of 97.0% for L, 96.1% for M, and 98.9% for S (**Supplemental Figure S5**). In contrast, the mean pairwise nucleotide identities to the SHAV-EU1 clade were 87.7% for L, 86.6% for M, and 95.8% for S (**Supplemental Figure S5**).

**Figure 2:**
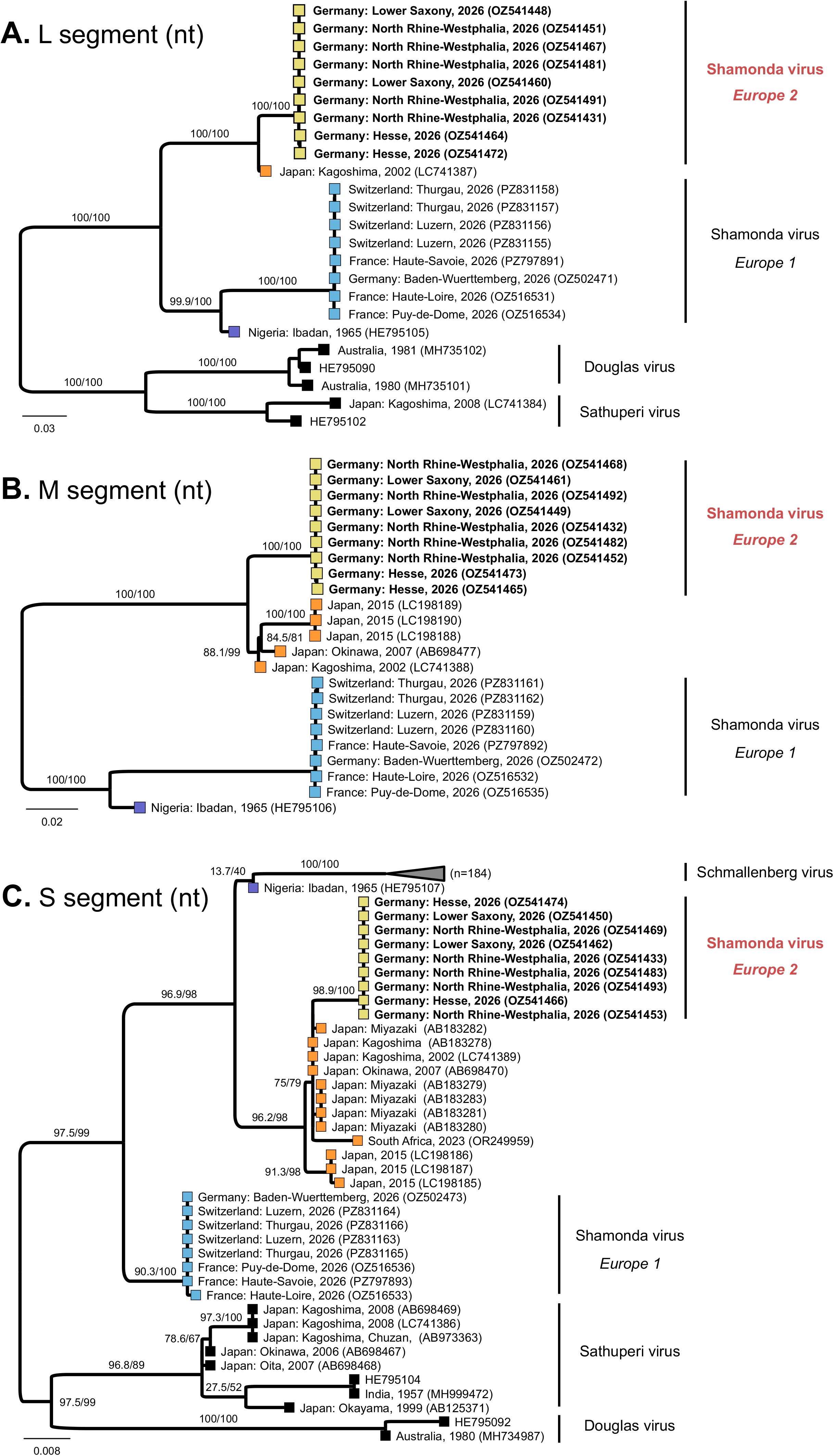
Maximum-likelihood phylogenetic analysis of the two newly detected Shamonda virus clades along with selected orthobunyaviruses. Coding sequences of the L protein (**A**.), the glycoprotein precursor (**B**.) and the nucleocapsid protein (**C**.) were extracted and aligned using MUSCLE (v5.1). The alignment was partitioned according to the first, second, and third codon positions to account for differences in evolutionary constraints and substitution rates among codon positions. Phylogenetic inference was performed with IQ-TREE (v3.1.3), using ModelFinder with partition merging (-m MFP+MERGE) to select the best-fitting substitution models and optimize the partitioning scheme. The maximum-likelihood tree search was performed using exhaustive nearest-neighbor interchange optimization (-allnni). Branch support was assessed using 10,000 SH-aLRT and 10,000 ultrafast bootstrap replicates, with additional nearest-neighbour interchange optimization of bootstrap trees (-bnni). Shamonda virus sequences are coloured according to their proposed clades. The newly detected Shamonda virus clade Europe 2 is highlighted in bold and red text. Branch support is shown for relevant nodes as SH-aLRT support (%) / ultrafast bootstrap support (%).

## Discussion

Here, we report the first detection and genomic characterization of a second clade of SHAV in Europe following the incursion of the SHAV-EU1 clade in Switzerland, France and southern Germany (1, 2). In order to facilitate the attribution of strains, we provisionally designated the clades as Shamonda virus Europe 1 (SHAV-EU1) for the variant circulating in southern Germany, Switzerland and France and Shamonda virus Europe 2 (SHAV-EU2) for the newly identified clade from northwestern Germany. Notably, both clades were detected during the same year but initially occurred in geographically separated areas, with the previously described clade SHAV-EU1 predominating in the south (1) and the clade reported here being identified in the northwest of Germany (**Figure 1**). The marked geographic separation of the initial detections, together with the genetic distinction between the two clades (**Figure 2**), indicates two independent introductions. The subsequent simultaneous detection of both clades in central Germany shows that their distribution ranges overlap and may reflect ongoing massive spread following their independent introductions. Importantly, the related SBV is present enzootically in the same regions, circulating in wave-like patterns, with years of high case numbers followed by periods of only sporadic detections (6-9). This situation also poses diagnostic challenges, as SBV must now be differentiated from two distinct SHAV variants, which in turn must be distinguished from each other. In addition, the host species affected by the two variants need to be considered separately. To date, samples positive for SHAV-EU2 have originated exclusively from cattle, whereas SHAV-EU1 has also been detected in horses presenting with central nervous disease (10). From a veterinary public health perspective, this further complicates the epidemiological picture and highlights the need for a differentiated assessment of the two SHAV variants.

Importantly, both SHAV clades differ markedly from SBV in the M segment (<40% amino acid identity), which encodes the major surface glycoproteins (11). Consequently, substantial cross-reactivity of the principal neutralizing immunogens is not expected, and pre-existing SBV immunity does not appear to have impeded the spread of SHAV. In contrast, SHAV-EU1 and SHAV-EU2 share more than 90% amino acid identity in the M-segment-encoded proteins, suggesting a substantial degree of antigenic cross-reactivity between the two variants. Experimental confirmation of this assumption, particularly by cross-neutralization studies, should therefore be considered a high priority.

As viruses of the Simbu serogroup possess tripartite genomes (11), co-circulation of several members of this virus group provides opportunities for reassortment if compatible viruses infect the same mammalian host or insect vector (11). Theoretically, three distinct viruses belonging to the species *Orthobunyavirus schmallenbergense* can generate 27 different segment combinations through reassortment. Excluding the three parental genotypes, this results in 24 potential reassortant genotypes. These reassortments may generate genomic constellations with altered biological properties and have to be monitored very closely.

In summary, the independent emergence of two distinct SHAV clades in Europe, followed by their rapid geographical spread, is a significant development with major implications for diagnostics, surveillance, animal health and risk evaluation. Of particular concern are the potential consequences of foetal infection, which are largely unknown and may only become apparent long afterwards through reproductive losses or congenital disease in affected animal species. This creates an additional challenge when it comes to assessing the full impact of the current epizootic situation. Furthermore, the introduction routes of both SHAV clades into Europe are still unknown and require further investigation to better understand their origins, dispersal, and epidemiological dynamics. Identifying these pathways will also be essential for determining whether targeted preventive or control measures can be developed to limit future introductions and spread.

## Statements

### Data availability

The sequences are available from the INSDC (International Nucleotide Sequence Database Collaboration, http://www.insdc.org) databases under project no. PRJEB125115.

### Ethical statement

The samples were taken by veterinarians in the context of health monitoring and as part of their clinical practice, no permissions were needed to collect these specimens. **Use of artificial intelligence tools** DeepL (DeepL SE, Cologne, Germany) and ChatGPT (OpenAI, San Francisco, CA, USA; GPT-5.6) were used solely for language editing and to improve the readability and linguistic clarity of the manuscript. The tools were not used for data analysis, interpretation of results, or generation of scientific conclusions. All generated or revised text was critically reviewed and approved by the authors, who take full responsibility for the final content.

## Acknowledgements

We thank Bianka Hillmann, Doreen, Schulz, Patrick Zitzow and Anja Landmesser-Zitzow for excellent technical assistance.

## References

1. Wernike K, Hoffmann B, Link EK, Rotheneder S, Eshak M, Pfaff F, et al. It happened again - emergence of a Shamonda-like orthobunyavirus in Central Europe, 2026. bioRxiv. 2026.

2. Kelleci M, Fusade-Boyer M, Chretien D, Durand E, Mircovich M, Secula A, et al. Detection of a novel Shamonda Orthobunyavirus in dairy cattle, France, June 2026. Euro Surveill. 2026;31(33).

3. Golender N, Bumbarov VY, Erster O, Beer M, Khinich Y, Wernike K. Development and validation of a universal S-segment-based real-time RT-PCR assay for the detection of Simbu serogroup viruses. J Virol Methods. 2018;261:80–5.

4. Bilk S, Schulze C, Fischer M, Beer M, Hlinak A, Hoffmann B. Organ distribution of Schmallenberg virus RNA in malformed newborns. Vet. Microbiol. 2012;159(1-2):236–8.

5. Fischer M, Schirrmeier H, Wernike K, Wegelt A, Beer M, Hoffmann B. Development of a pan-Simbu real-time reverse transcriptase PCR for the detection of Simbu serogroup viruses and comparison with SBV diagnostic PCR systems. Virol. J. 2013;10(1):327.

6. Larska M. Schmallenberg virus: a cyclical problem. Vet Rec. 2018;183(22):688–9.

7. Zeiske S, Kampen H, Sick F, Dähn O, Voigt A, Heuser E, et al. Monitoring of Schmallenberg virus, bluetongue virus and epizootic haemorrhagic disease virus in biting midges in Germany 2019-2023. Parasites & vectors. 2025;18(1):262.

8. Bayrou C, Lesenfants C, Paternostre J, Volpe R, Moula N, Coupeau D, et al. Schmallenberg virus, cyclical reemergence in the core region: A seroepidemiologic study in wild cervids, Belgium, 2012-2017. Transbound. Emerg. Dis. 2022;69(3):1625–33.

9. Wernike K, Fischer L, Twietmeyer S, Beer M. Extensive Schmallenberg virus circulation in Germany, 2023. Vet. Res. 2024;55(1):134.

10. Institute of Virology and Immunology. Shamonda virus in Switzerland. Online available: https://www.ivi.admin.ch/en/neues-orthobunyavirus-der-simbu-serogruppe-bei-rindern-entdeckte, last accessed 04 September 2026. 2026.

11. Elliott RM. Orthobunyaviruses: recent genetic and structural insights. Nat. Rev. Microbiol. 2014;12(10):673–85.

