## Supplemental Material for "Two incursions, two viruses: Emergence of a second novel Shamonda virus clade in Central Europe, 2026"

for

#### Supplemental Material and Methods

##### Metagenomic sequencing

RNA was extracted from plasma samples using the RNAdvance Tissue Kit (Beckman Coulter) on a KingFisher Duo Prime platform (Thermo Fisher Scientific). For sequencing, the RNA was further processed as previously described [1]. In addition, after size selection, the libraries were amplified using AccuPrime Pfx DNA Polymerase and subsequently size selected again. Thereafter, libraries were quantified as described [1]. Pooled libraries were loaded onto Ion 530 chips using an Ion Chef instrument and sequenced on an Ion S5 XL platform in 400-bp mode. The obtained datasets were assembled with Newbler (v3.0; Roche/454) and for nine samples a segmented viral genome with similarity to orthobunyaviruses was obtained. The resulting complete coding sequences are available from the INSDC (International Nucleotide Sequence Database Collaboration, http://www.insdc.org) databases under project no. PRJEB125115.

#### Phylogenetic analysis

Phylogenetic analyses were performed separately for all three genome segments.

Initially, for a broader phylogenetic classification appropriate reference sequences (RefSeq) of the genus *Orthobunyavirus* were retrieved from GenBank and viruses belonging to the Simbu, Bunyamwera, California, Leanyer and Sedlec serogroups were selected. The amino acid sequences were aligned using MUSCLE (v5.1; [2]) and Maximum-likelihood trees were inferred with IQ-TREE (v3.1.3; [3]) using ModelFinder (-m MFP; [4]) to select the best-fitting amino acid substitution model. Tree searches were performed using exhaustive nearest-neighbor interchange optimization (-allnni). Branch support was assessed using 10,000 ultrafast bootstrap replicates [5] and 10,000 SH-aLRT replicates. Bootstrap trees were additionally optimized using nearest-neighbor interchange (-bnni).

For the analyses of nucleotide sequences, reference sequences from the species *Orthobunyavirus schmallenbergense* (Douglas virus, Sathuperi virus, Schmallenberg virus and Shamonda virus) were retrieved from GenBank. Subsequently, the coding sequences (CDSs) were first identified using "Find ORF" function implemented in Geneious Prime (v2025.1.3) and subsequently extracted. The extracted CDSs were aligned using MUSCLE (v5.1) and Maximum-likelihood phylogenetic trees were inferred with IQ-TREE (v3.1.3; [3]) using a partitioned analysis [6]. In detail, the alignments were partitioned by codon position (first, second, and third positions) to account for the possible different evolutionary constraints and substitution rates among codon positions. ModelFinder was used to determine the best-fitting substitution model for each partition and to optimize the partitioning scheme (-m MFP+MERGE; [4]). Branch support statistics and NNI optimization was done as described above.

### Supplemental Figures

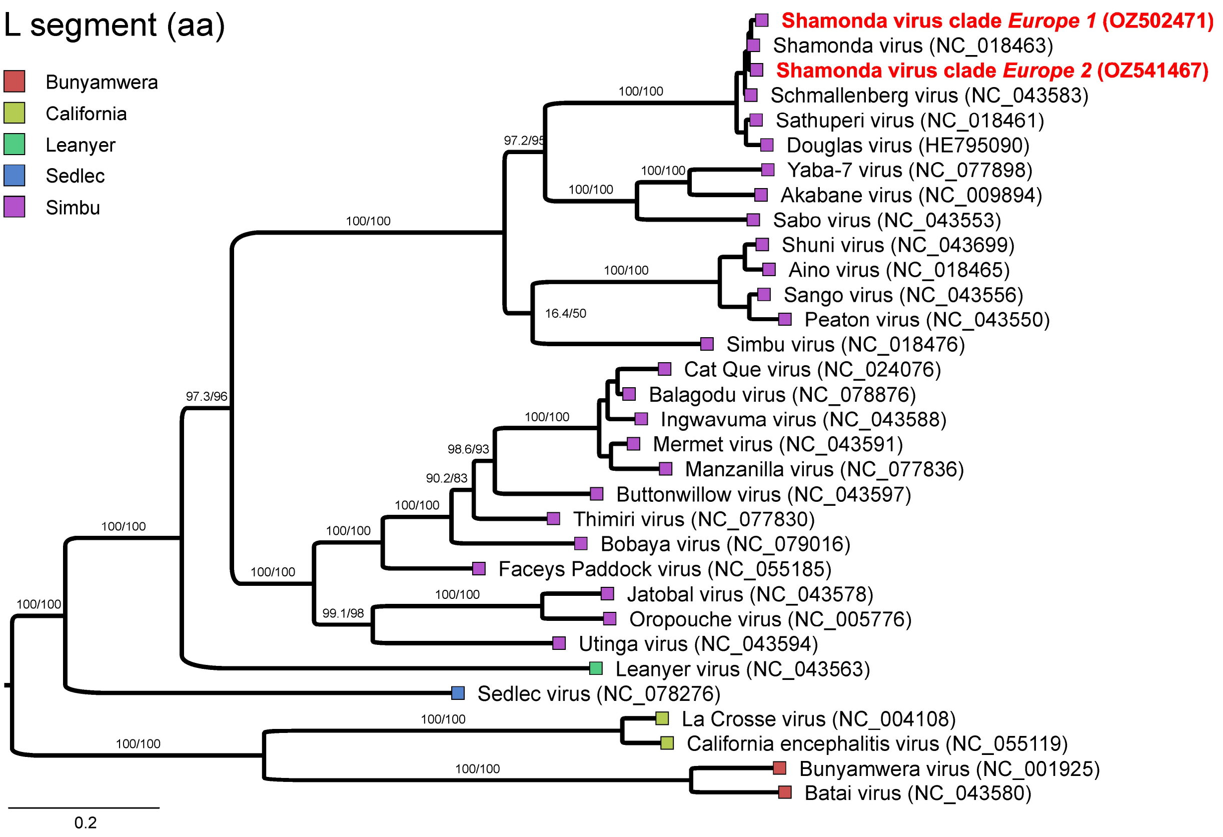


**Supplemental Figure S1: Maximum-likelihood phylogenetic analysis of the L segment of selected orthobunyaviruses.** Coding sequences of the L protein were extracted, translated, and aligned using MUSCLE (v5.1). Phylogenetic inference was performed with IQ-TREE (v3.1.3), with the optimal amino acid substitution model selected using ModelFinder (-m MFP). The maximum-likelihood tree search was performed using exhaustive nearest-neighbor interchange optimization (-allnni). Branch support was assessed using 10,000 SH-aLRT and 10,000 ultrafast bootstrap replicates, with additional nearest-neighbour interchange optimization of bootstrap trees (-bnni). Viruses are coloured according to their serogroup. The two newly detected Shamonda virus clades, Europe 1 and Europe 2, are highlighted in bold red text. Branch support is shown for relevant nodes as SH-aLRT support (%) / ultrafast bootstrap support (%).

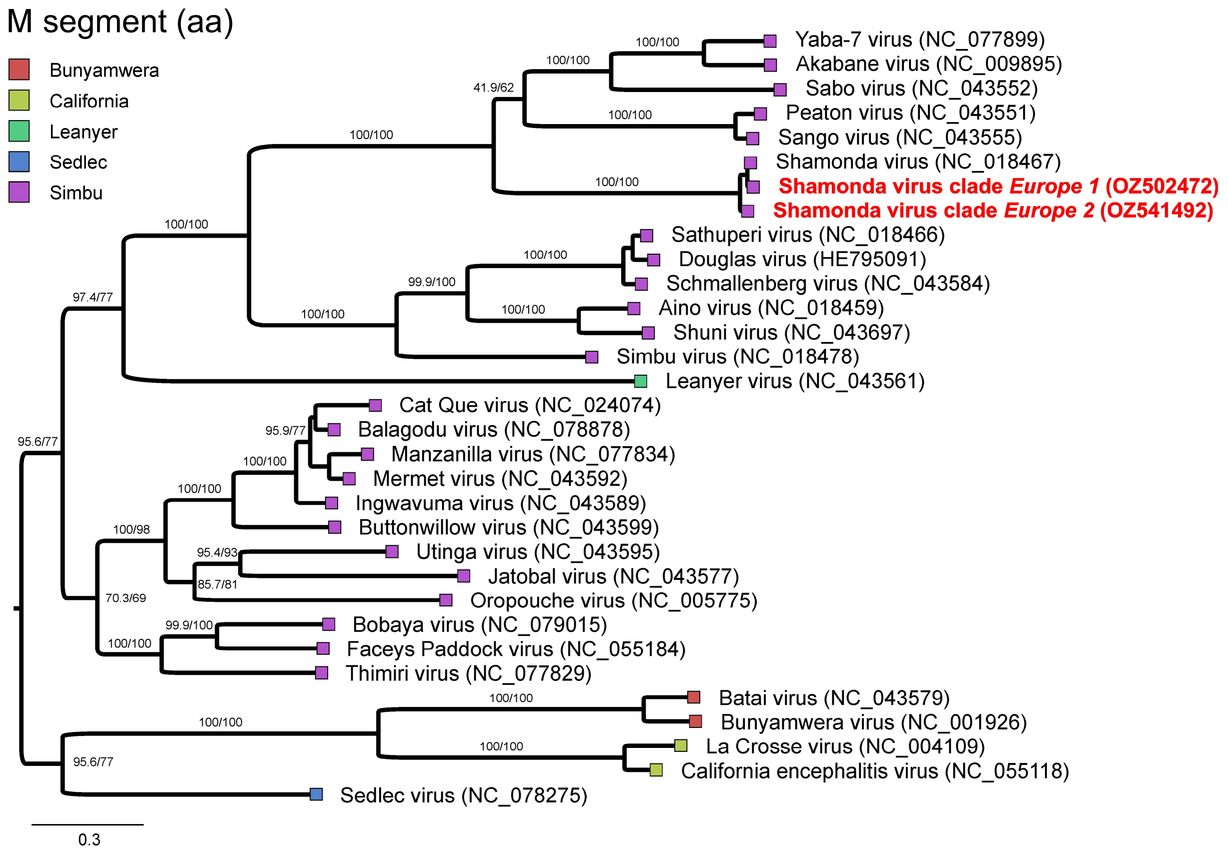


**Supplemental Figure S2: Maximum-likelihood phylogenetic analysis of the M segment of selected orthobunyaviruses.** Coding sequences of the glycoprotein precursor were extracted, translated, and aligned using MUSCLE (v5.1). Phylogenetic inference was performed with IQ-TREE (v3.1.3), with the optimal amino acid substitution model selected using ModelFinder (-m MFP). The maximum-likelihood tree search was performed using exhaustive nearest-neighbor interchange optimization (-allnni). Branch support was assessed using 10,000 SH-aLRT and 10,000 ultrafast bootstrap replicates, with additional nearest-neighbour interchange optimization of bootstrap trees (-bnni). Viruses are coloured according to their serogroup. The two newly detected Shamonda virus clades, Europe 1 and Europe 2, are highlighted in bold red text. Branch support is shown for relevant nodes as SH-aLRT support (%) / ultrafast bootstrap support (%).

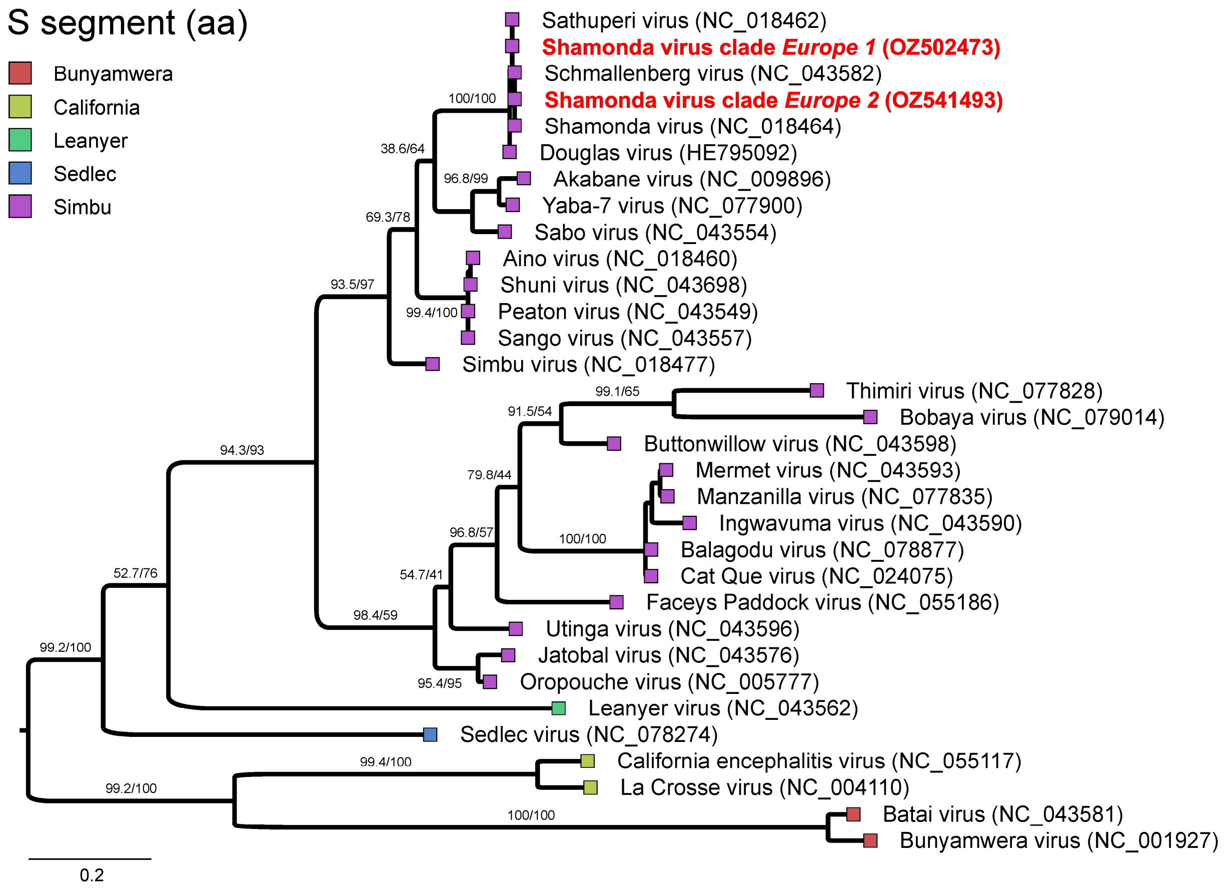


**Supplemental Figure S3: Maximum-likelihood phylogenetic analysis of the S segment of selected orthobunyaviruses.** Coding sequences of the nucleocapsid protein were extracted, translated, and aligned using MUSCLE (v5.1). Phylogenetic inference was performed with IQ-TREE (v3.1.3), with the optimal amino acid substitution model selected using ModelFinder (-m MFP). The maximum-likelihood tree search was performed using exhaustive nearest-neighbor interchange optimization (-allnni). Branch support was assessed using 10,000 SH-aLRT and 10,000 ultrafast bootstrap replicates, with additional nearest-neighbour interchange optimization of bootstrap trees (-bnni). Viruses are coloured according to their serogroup. The two newly detected Shamonda virus clades, Europe 1 and Europe 2, are highlighted in bold red text. Branch support is shown for relevant nodes as SH-aLRT support (%) / ultrafast bootstrap support (%).

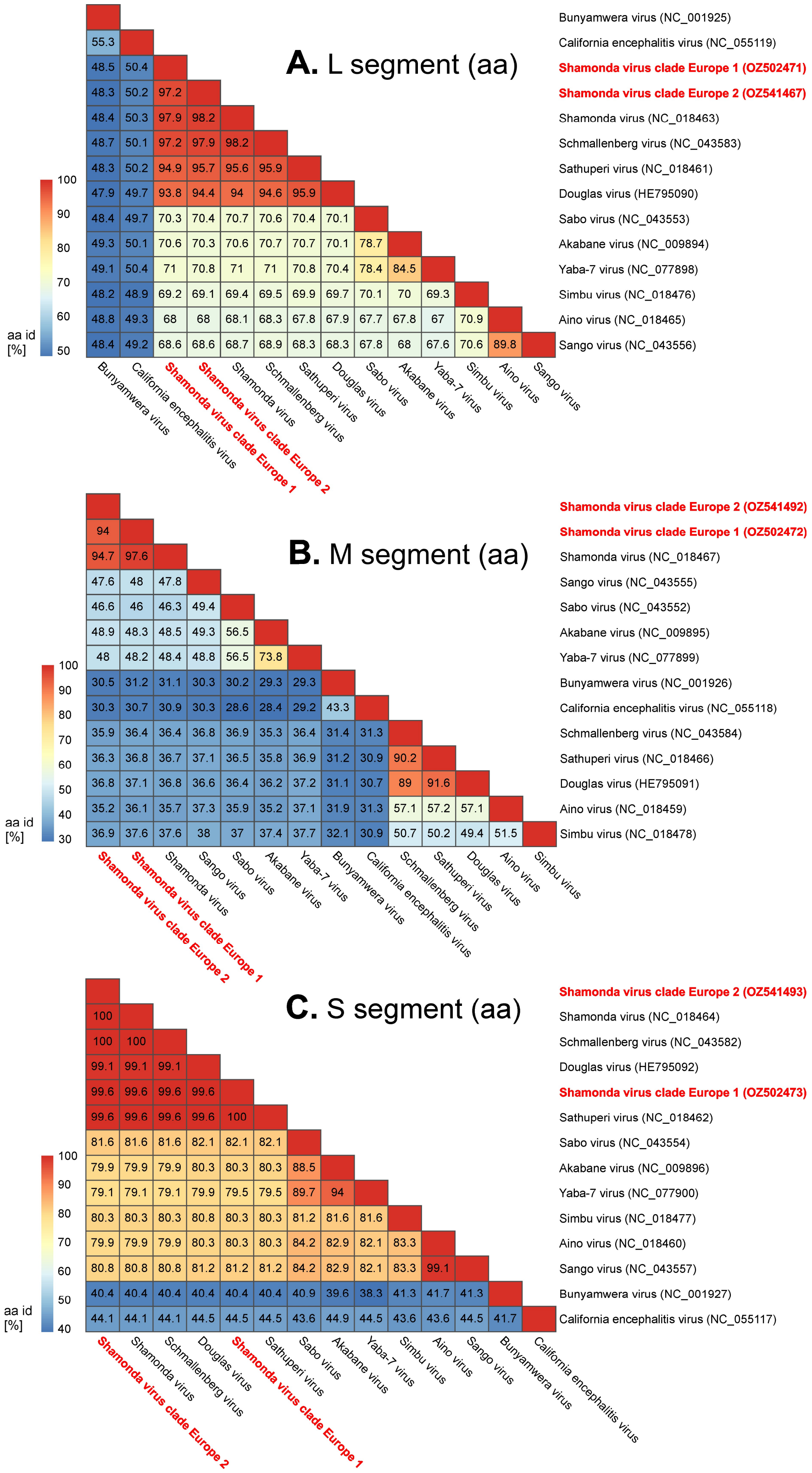


**Supplemental Figure S4: Pairwise amino acid identity of selected orthobunyaviruses.** Coding sequences of the L protein (**A.**), the glycoprotein precursor (**B.**) and the nucleocapsid protein (**C.**) were extracted, translated, and aligned using MUSCLE (v5.1). The pairwise amino acid identity was deduced and plotted. The two newly detected Shamonda virus clades, Europe 1 and Europe 2, are highlighted in bold red text.

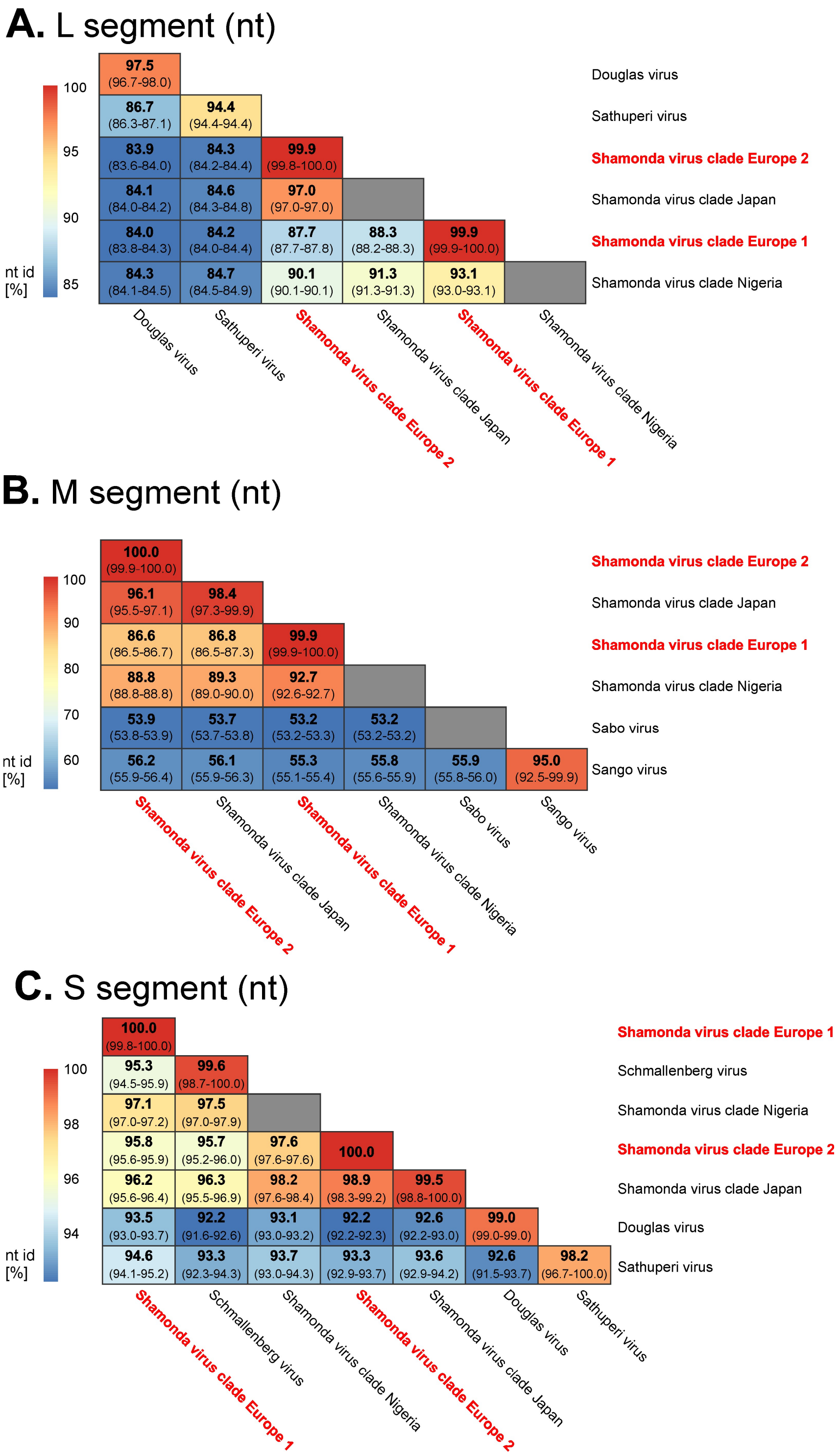


**Supplemental Figure S5: Pairwise nucleotide identity of selected orthobunyaviruses.** Coding sequences of the L protein (**A.**), the glycoprotein precursor (**B.**) and nucleocapsid protein (**C.**) were extracted and aligned using MUSCLE (v5.1). Pairwise nucleotide identities were calculated for all included sequences and summarized for each virus pair as the mean identity, with the corresponding minimum–maximum range shown below. The two newly detected Shamonda virus clades, Europe 1 and Europe 2, are highlighted in bold red text.

### Supplemental Table

**Supplemental Table S1: Data on samples sent to the German National Reference Laboratory for Schmallenberg virus from animals suspected of having a Shamonda virus infection.** Origin, sample material, date of sample receipt and RT-qPCR results are given for samples that tested positive by a pan-Simbuvirus RT-qPCR. Samples that were investigated by metagenomic sequencing are marked by an asterisk. BW - Baden-Württemberg, BY - Bavaria, HE - Hesse, NI - Lower Saxony, NW - North Rhine-Westphalia, RP - Rhineland-Palatinate

| Sample ID | Origin | Sample material | Date | Pan-Simbu RT-qPCR | S-segment-based SBV RT-qPCR | L-segment-based SBV RT-qPCR |
|---|---|---|---|---|---|---|
| 2026BVD09542 | Germany, NW | serum | 18-08-2026 | 27.7 / positive | 28.7 / positive | negative |
| 2026BVD09543 | Germany, NW | serum | 18-08-2026 | 26.7 / positive | 28.5 / positive | negative |
| 2026BVD09544 | Germany, NW | serum | 18-08-2026 | 24.9 / positive | 26.6 / positive | negative |
| 2026BVD09545 | Germany, NW | serum | 18-08-2026 | 28.8 / positive | 30.5 / positive | negative |
| 2026BVD09546 | Germany, NW | serum | 18-08-2026 | 32.3 / positive | 32.8 / positive | negative |
| 2026BVD09547 | Germany, NW | serum | 18-08-2026 | 33.9 / positive | 34.2 / positive | negative |
| 2026BVD09548 | Germany, NW | serum | 18-08-2026 | 36.2 / positive | 39.7 / positive | negative |
| 2026BVD09699* | Germany, NW | plasma | 19-08-2026 | 23.7 / positive | 27.1 / positive | negative |
| 2026BVD09701* | Germany, NW | plasma | 19-08-2026 | 22.8 / positive | 26.3 / positive | negative |
| 2026BVD09702 | Germany, NW | plasma | 19-08-2026 | 24.9 / positive | 28.1 / positive | negative |
| 2026BVD09703 | Germany, NW | plasma | 19-08-2026 | 23.9 / positive | 26.5 / positive | negative |
| 2026BVD09711 | Germany, NW | plasma | 19-08-2026 | 33.3 / positive | 35.1 / positive | negative |
| 2026BVD09712* | Germany, NW | plasma | 19-08-2026 | 23.3 / positive | 26.4 / positive | negative |
| 2026BVD09714 | Germany, NW | plasma | 19-08-2026 | 24.1 / positive | 26.6 / positive | negative |
| 2026BVD09715 | Germany, NW | plasma | 19-08-2026 | 25.4 / positive | 28.2 / positive | negative |
| 2026BVD09716 | Germany, NW | plasma | 19-08-2026 | 26.2 / positive | 28.0 / positive | negative |
| 2026BVD09717 | Germany, NW | plasma | 19-08-2026 | 27.8 / positive | 29.6 / positive | negative |
| 2026BVD09722 | Germany, NW | plasma | 19-08-2026 | 24.3 / positive | 27.2 / positive | negative |
| 2026BVD09724 | Germany, NW | plasma | 19-08-2026 | 34.8 / positive | 35.2 / positive | negative |
| 2026BVD09725 | Germany, NW | plasma | 19-08-2026 | 25.3 / positive | 28.5 / positive | negative |
| 2026BVD09726 | Germany, NW | plasma | 19-08-2026 | 24.7 / positive | 27.4 / positive | negative |
| 2026BVD09727 | Germany, NW | plasma | 19-08-2026 | 35.2 / positive | 39.6 / positive | negative |
| 2026BVD09728 | Germany, NW | plasma | 19-08-2026 | 30.2 / positive | 31.7 / positive | negative |
| 2026BVD09729 | Germany, NW | serum | 19-08-2026 | 23.7 / positive | 26.1 / positive | negative |

| Sample ID | Origin | Sample material | Date | Pan-Simbu RT-qPCR | S-segment-based SBV RT-qPCR | L-segment-based SBV RT-qPCR |
|---|---|---|---|---|---|---|
| 2026BVD09753 | Germany, NW | EDTA blood | 20-08-2026 | 23.9 / positive | 26.3 / positive | negative |
| 2026BVD09754 | Germany, NW | EDTA blood | 20-08-2026 | 29.8 / positive | 28.9 / positive | negative |
| 2026BVD09755 | Germany, NW | EDTA blood | 20-08-2026 | 26.4 / positive | 26.1 / positive | negative |
| 2026BVD09756 | Germany, NW | EDTA blood | 20-08-2026 | 25.8 / positive | 29.2 / positive | negative |
| 2026BVD09757 | Germany, NW | EDTA blood | 20-08-2026 | 26.9 / positive | 33.2 / positive | negative |
| 2026BVD09758 | Germany, NW | serum | 20-08-2026 | 26.7 / positive | 28.4 / positive | negative |
| 2026BVD09759 | Germany, NW | serum | 20-08-2026 | 24.2 / positive | 26.2 / positive | negative |
| 2026BVD09760 | Germany, NW | serum | 20-08-2026 | 24.3 / positive | 26.5 / positive | negative |
| 2026BVD09761 | Germany, NW | serum | 20-08-2026 | 26.5 / positive | 27.6 / positive | negative |
| 2026BVD09768 | Germany, NW | EDTA blood | 20-08-2026 | 24.5 / positive | 26.2 / positive | negative |
| 2026BVD09769 | Germany, NW | EDTA blood | 20-08-2026 | 23.3 / positive | 25.3 / positive | negative |
| 2026BVD09770 | Germany, NW | EDTA blood | 20-08-2026 | 24.1 / positive | 26.5 / positive | negative |
| 2026BVD09771* | Germany, NW | EDTA blood | 20-08-2026 | 21.7 / positive | 24.6 / positive | negative |
| 2026BVD09772 | Germany, NW | EDTA blood | 20-08-2026 | 25.7 / positive | 27.3 / positive | negative |
| 2026BVD09774* | Germany, NW | EDTA blood | 20-08-2026 | 21.6 / positive | 24.1 / positive | negative |
| 2026BVD09775 | Germany, NW | EDTA blood | 20-08-2026 | 30.8 / positive | 32.9 / positive | negative |
| 2026BVD09776 | Germany, NW | EDTA blood | 20-08-2026 | 29.5 / positive | 30.8 / positive | negative |
| 2026BVD09780 | Germany, NW | EDTA blood | 20-08-2026 | 24.5 / positive | 27.3 / positive | negative |
| 2026BVD09549 | Germany, NI | EDTA blood | 18-08-2026 | 30.6 / positive | 30.3 / positive | negative |
| 2026BVD09818 | Germany, NI | EDTA blood | 20-08-2026 | 25.0 / positive | 26.3 / positive | negative |
| 2026BVD09819 | Germany, NI | EDTA blood | 20-08-2026 | 20.3 / positive | 23.9 / positive | negative |
| 2026BVD09820 | Germany, NI | EDTA blood | 20-08-2026 | 25.2 / positive | 26.1 / positive | negative |
| 2026BVD09821* | Germany, NI | EDTA blood | 20-08-2026 | 23.0 / positive | 24.8 / positive | negative |
| 2026BVD09822 | Germany, NI | EDTA blood | 20-08-2026 | 24.4 / positive | 26.6 / positive | negative |
| 2026BVD09823 | Germany, NI | EDTA blood | 20-08-2026 | 32.5 / positive | 33.3 / positive | negative |
| 2026BVD09824 | Germany, NI | EDTA blood | 20-08-2026 | 25.6 / positive | 27.3 / positive | negative |
| 2026BVD09825 | Germany, NI | EDTA blood | 20-08-2026 | 35.3 / positive | 35.5 / positive | negative |
| 2026BVD09826 | Germany, NI | EDTA blood | 20-08-2026 | 36.3 / positive | 37.7 / positive | negative |
| 2026BVD09828 | Germany, NI | EDTA blood | 20-08-2026 | 29.3 / positive | 29.4 / positive | negative |
| 2026BVD09829* | Germany, NI | EDTA blood | 20-08-2026 | 23.3 / positive | 25.2 / positive | negative |
| 2026BVD09830 | Germany, NI | EDTA blood | 20-08-2026 | 27.5 / positive | 28.9 / positive | negative |
| 2026BVD09831 | Germany, NI | EDTA blood | 20-08-2026 | 33.2 / positive | 34.0 / positive | negative |
| 2026BVD09210 | Germany, HE | EDTA blood | 13-08-2026 | 22.1 / positive | negative | 22.7 / positive |
| 2026BVD09784 | Germany, HE | EDTA blood | 20-08-2026 | 26.0 / positive | 27.6 / positive | negative |

| Sample ID | Origin | Sample material | Date | Pan-Simbu RT-qPCR | S-segment-based SBV RT-qPCR | L-segment-based SBV RT-qPCR |
|---|---|---|---|---|---|---|
| 2026BVD09785 | Germany, HE | EDTA blood | 20-08-2026 | 31.1 / positive | 31.9 / positive | negative |
| 2026BVD09786 | Germany, HE | EDTA blood | 20-08-2026 | 21.2 / positive | negative | 23.8 / positive |
| 2026BVD09788 | Germany, HE | EDTA blood | 20-08-2026 | 24.7 / positive | negative | 27.2 / positive |
| 2026BVD09790 | Germany, HE | EDTA blood | 20-08-2026 | 26.6 / positive | negative | 27.8 / positive |
| 2026BVD09791 | Germany, HE | EDTA blood | 20-08-2026 | 21.6 / positive | negative | 23.9 / positive |
| 2026BVD09792 | Germany, HE | EDTA blood | 20-08-2026 | 26.4 / positive | negative | 28.2 / positive |
| 2026BVD09793* | Germany, HE | EDTA blood | 20-08-2026 | 22.4 / positive | 24.6 / positive | negative |
| 2026BVD09794 | Germany, HE | EDTA blood | 20-08-2026 | 29.6 / positive | 30.4 / positive | negative |
| 2026BVD09795 | Germany, HE | EDTA blood | 20-08-2026 | 29.0 / positive | 30.5 / positive | negative |
| 2026BVD09796 | Germany, HE | EDTA blood | 20-08-2026 | 37.8 / positive | 36.5 / positive | negative |
| 2026BVD09797* | Germany, HE | EDTA blood | 20-08-2026 | 24.4 / positive | 26.3 / positive | negative |
| 2026BVD09798 | Germany, HE | serum | 20-08-2026 | 20.8 / positive | negative | 23.0 / positive |
| 2026BVD09801 | Germany, HE | EDTA blood | 20-08-2026 | 20.9 / positive | negative | 23.8 / positive |
| 2026BVD09802 | Germany, HE | EDTA blood | 20-08-2026 | 25.5 / positive | negative | 27.4 / positive |
| 2026BVD09803 | Germany, HE | EDTA blood | 20-08-2026 | 27.7 / positive | 29.0 / positive | negative |
| 2026BVD09804 | Germany, HE | EDTA blood | 20-08-2026 | 22.3 / positive | negative | 24.7 / positive |
| 2026BVD09805 | Germany, HE | EDTA blood | 20-08-2026 | 25.3 / positive | 27.1 / positive | negative |
| 2026BVD09806 | Germany, HE | EDTA blood | 20-08-2026 | 25.2 / positive | negative | 28.2 / positive |
| 2026BVD09809 | Germany, HE | EDTA blood | 20-08-2026 | 34.4 / positive | 34.0 / positive | negative |
| 2026BVD09731 | Germany, BW | EDTA blood | 19-08-2026 | 32.7 / positive | negative | 32.9 / positive |
| 2026BVD09732 | Germany, BW | EDTA blood | 19-08-2026 | 22.2 / positive | negative | 23.9 / positive |
| 2026BVD09733 | Germany, BW | serum | 19-08-2026 | 29.2 / positive | negative | 30.3 / positive |
| 2026BVD09734 | Germany, BW | serum | 19-08-2026 | 23.0 / positive | negative | 24.9 / positive |
| 2026BVD09735 | Germany, BW | EDTA blood | 19-08-2026 | 21.1 / positive | negative | 23.5 / positive |
| 2026BVD09737 | Germany, BW | EDTA blood | 19-08-2026 | 22.1 / positive | negative | 23.6 / positive |
| 2026BVD09739 | Germany, BW | serum | 19-08-2026 | 22.2 / positive | negative | 24.0 / positive |
| 2026BVD09740 | Germany, BW | EDTA blood | 19-08-2026 | 21.5 / positive | negative | 23.1 / positive |
| 2026BVD09741 | Germany, BW | EDTA blood | 19-08-2026 | 24.8 / positive | negative | 27.5 / positive |
| 2026BVD09742 | Germany, BW | serum | 19-08-2026 | 20.2 / positive | negative | 22.7 / positive |
| 2026BVD09743 | Germany, BW | serum | 19-08-2026 | 23.1 / positive | negative | 25.6 / positive |
| 2026BVD09744 | Germany, BW | EDTA blood | 19-08-2026 | 20.5 / positive | negative | 23.4 / positive |
| 2026BVD09745 | Germany, BW | EDTA blood | 19-08-2026 | 19.3 / positive | negative | 21.8 / positive |
| 2026BVD09746 | Germany, BW | serum | 19-08-2026 | 22.4 / positive | negative | 25.0 / positive |
| 2026BVD09747 | Germany, BW | serum | 19-08-2026 | 18.5 / positive | negative | 22.3 / positive |

| Sample ID | Origin | Sample material | Date | Pan-Simbu RT-qPCR | S-segment-based SBV RT-qPCR | L-segment-based SBV RT-qPCR |
|---|---|---|---|---|---|---|
| 2026BVD09260 | Germany, BY | serum | 14-08-2026 | 19.6 / positive | negative | 21.5 / positive |
| 2026BVD09261 | Germany, BY | serum | 14-08-2026 | 24.9 / positive | negative | 27.0 / positive |
| 2026BVD09262 | Germany, BY | serum | 14-08-2026 | 37.2 / positive | negative | 39.7 / positive |
| 2026BVD09263 | Germany, BY | serum | 14-08-2026 | 33.0 / positive | negative | 35.9 / positive |
| 2026BVD09264 | Germany, BY | serum | 14-08-2026 | 34.4 / positive | negative | 37.0 / positive |
| 2026BVD09265 | Germany, BY | serum | 14-08-2026 | 19.8 / positive | negative | 20.7 / positive |
| 2026BVD09266 | Germany, BY | serum | 14-08-2026 | 24.1 / positive | negative | 25.1 / positive |
| 2026BVD09267 | Germany, BY | serum | 14-08-2026 | 35.3 / positive | negative | 38.6 / positive |
| 2026BVD09269 | Germany, BY | serum | 14-08-2026 | 25.1 / positive | negative | 26.4 / positive |
| 2026BVD09270 | Germany, BY | serum | 14-08-2026 | 22.3 / positive | negative | 23.1 / positive |
| 2026BVD07609 | Germany, RP | EDTA blood | 06-08-2026 | 26.4 / positive | negative | 26.5 / positive |
| 2026BVD07611 | Germany, RP | EDTA blood | 06-08-2026 | 24.6 / positive | negative | 24.2 / positive |
| 2026BVD07613 | Germany, RP | EDTA blood | 06-08-2026 | 25.1 / positive | negative | 25.4 / positive |
| 2026BVD07614 | Germany, RP | EDTA blood | 06-08-2026 | 23.7 / positive | negative | 23.4 / positive |
| 2026BVD07615 | Germany, RP | EDTA blood | 06-08-2026 | 33.2 / positive | negative | 31.9 / positive |
| 2026BVD07619 | Germany, RP | EDTA blood | 06-08-2026 | 29.6 / positive | negative | 28.5 / positive |
| 2026BVD08010 | Germany, RP | EDTA blood | 06-08-2026 | 22.4 / positive | negative | 22.7 / positive |
| 2026BVD08011 | Germany, RP | EDTA blood | 06-08-2026 | 24.4 / positive | negative | 24.6 / positive |
| 2026BVD08012 | Germany, RP | EDTA blood | 06-08-2026 | 23.8 / positive | negative | 22.1 / positive |
| 2026BVD08013 | Germany, RP | EDTA blood | 06-08-2026 | 24.3 / positive | negative | 23.5 / positive |
| 2026BVD08015 | Germany, RP | EDTA blood | 06-08-2026 | 23.3 / positive | negative | 24.0 / positive |
| 2026BVD08016 | Germany, RP | EDTA blood | 06-08-2026 | 26.1 / positive | negative | 25.3 / positive |
| 2026BVD08017 | Germany, RP | EDTA blood | 06-08-2026 | 28.0 / positive | negative | 27.2 / positive |
| 2026BVD08018 | Germany, RP | EDTA blood | 06-08-2026 | 23.0 / positive | negative | 23.4 / positive |
| 2026BVD08019 | Germany, RP | EDTA blood | 06-08-2026 | 25.0 / positive | negative | 25.2 / positive |
| 2026BVD08020 | Germany, RP | EDTA blood | 06-08-2026 | 25.3 / positive | negative | 25.0 / positive |
| 2026BVD08021 | Germany, RP | EDTA blood | 06-08-2026 | 24.1 / positive | negative | 21.3 / positive |
| 2026BVD08022 | Germany, RP | EDTA blood | 06-08-2026 | 25.0 / positive | negative | 24.9 / positive |
| 2026BVD08023 | Germany, RP | EDTA blood | 06-08-2026 | 26.6 / positive | negative | 27.1 / positive |
| 2026BVD08024 | Germany, RP | EDTA blood | 06-08-2026 | 34.4 / positive | negative | 39.4 / positive |

92

#### Supplemental References


1. Eschbaumer M, Staubach C, Pfaff F, et al. Buffaloed in Brandenburg: Germany’s first Brush with Foot-and-Mouth Disease after four Decades of Freedom, **2026**.
2. Edgar RC. MUSCLE: multiple sequence alignment with high accuracy and high throughput. Nucleic Acids Res **2004**; 32:1792–7.
3. Wong TKF, Ly-Trong N, Ren H, et al. IQ-TREE 3: phylogenomic inference software using complex evolutionary models. Mol Biol Evol **2026**; 43.
4. Kalyaanamoorthy S, Minh BQ, Wong TKF, Haeseler A von, Jermiin LS. ModelFinder: fast model selection for accurate phylogenetic estimates. Nat Methods **2017**; 14:587–9.
5. Hoang DT, Chernomor O, Haeseler A von, Minh BQ, Le Vinh S. UFBoot2: Improving the Ultrafast Bootstrap Approximation. Mol Biol Evol **2018**; 35:518–22.
6. Chernomor O, Haeseler A von, Minh BQ. Terrace Aware Data Structure for Phylogenomic Inference from Supermatrices. Syst Biol **2016**; 65:997–1008.
